# InCytokine, an open-source software, reveals a TREM2 variant specific cytokine signature

**DOI:** 10.1101/2025.10.31.685741

**Authors:** Deepak Jha, Marco Ancona, Filip Oplt, Sonia L. Farmer, Martin Vagenknecht, Alejandro Vazquez-Otero, Illia Pradznyk, Jindrich Soukup, Rebecca S. Mathew, Vanessa Peterson, Danny A. Bitton

**Affiliations:** Data and Genome Sciences, Merck & Co., Inc., Boston, Massachusetts, USA; Applied Research and Innovation, MSD Czech Republic s.r.o., Prague, Czech Republic; Data Science and Scientific Informatics, MSD Czech Republic s.r.o., Prague, Czech Republic

**Keywords:** Cytokines profiling, protein array, Alzheimer’s Disease, TREM2, DPP4, Biomarker

## Abstract

Cytokine and chemokine profiling is central to understanding inflammatory processes and the mechanisms driving diverse diseases. We introduce InCytokine, an open-source tool for semiquantitative analysis of cytokine and chemokine data generated by protein array technologies. InCytokine features robust and modular image-processing workflows, including automated spot detection, template alignment, normalization, quality-control measures and quantitative intensity summarization to deliver consistent and reliable readouts from profiling assays. We evaluated InCytokine by profiling wild-type microglia, TREM2 knockout, and Alzheimer’s-associated *TREM2 R47H* variant cells in response of lipopolysaccharide and sulfatide exposure. Differential expression analysis revealed unique sulfatide-specific and genotype-specific cytokine signatures in TREM2 variants. We also report an intriguing modulation of DPP4 and a divergent expression pattern of ENA-78 in TREM2 variants in response to lipopolysaccharide and sulfatide treatment. Such distinct expression signatures raise the possibility that TREM2 variants may play a role in modulating inflammatory signaling relevant to cardio-metabolic and Alzheimer’s disease. These signatures were corroborated using transcriptional profiling of the same microglia cells, revealing also a good concordance between protein array and RNA sequencing technologies. Taken together, InCytokine is an interactive, user-friendly web application for rapid, reproducible, and scalable analysis of protein array data, proven to generate meaningful insights for drug and biomarker discovery campaigns in pharmaceutical settings.

## Background

Inflammation is a key driver of diverse diseases including cancer, neurodegeneration, and aging, mediated through coordinated interplay between innate and adaptive immune responses [1–5]. Cytokines and chemokines orchestrate proliferation, differentiation, migration, as well as the initiation, amplification and resolution of immune responses. Chemokines specifically direct leukocyte positioning in homeostasis and inflammation [6–8]. Consequently, cytokine/chemokine profiles are widely used as biomarkers for early disease detection, therapeutic monitoring and disease staging [9–11].

Alzheimer’s disease (AD), the leading cause of dementia (60–80% of cases), affected 6.2 million Americans aged ≥65 in 2021 and may approach 14 million by 2050. AD ranks among the top causes of death in the U.S. with mortality exceeding that of breast and prostate cancer combined [12]. Neuroinflammation is now recognized as a hallmark of AD, involving activation of microglia and astrocytes and release of cytokines and chemokines in response to infection, injury and amyloid/tau pathology [3, 13–15]. Chronic glial activation contributes to synaptic dysfunction, neuronal loss and cognitive decline. Microglia exhibit context dependent roles supporting synaptic pruning and debris clearance and, early in disease, aiding Aβ clearance, but later driving neurotoxicity via proinflammatory cytokines and reactive oxygen species [3, 13, 16, 17]. These context-dependent dynamics underscore the value of systematic cytokine and chemokine profiling to elucidate underlying disease mechanisms and to identify novel therapeutic targets amenable to targeted intervention.

Multiple platforms enable such profiling with distinct tradeoffs, as follows. ELISA offers high specificity but is largely single-plex [18]. Multiplex bead-based immunoassays (e.g. Luminex) and cytometric bead arrays increase throughput and conserve sample volume but require control of cross reactivity and standardized analysis [19, 20]. Electrochemiluminescence immunoassays extend dynamic range and sensitivity but often depend on proprietary panels [21]. Proximity extension assays (Olink) and aptamer-based proteomics (SomaScan) support high-plex profiling with strong analytical sensitivity yet demand rigorous normalization and cross platform calibration [22–24]. Ultrasensitive digital immunoassays (e.g. Simoa) reach sub-femtomolar detection, enabling low abundance cytokine quantification with added analytical complexity [25]. Complementary modalities include flow cytometry for intracellular cytokines at single cell resolution (workflow intensive) and mass spectrometry proteomics for unbiased surveys that remain challenged for very low abundance targets in plasma/CSF [19, 26–28]. Membrane and slide based antibody arrays provide rapid, semi-quantitative chemiluminescent readouts across many analytes, but limited dynamic range, predefined panels and subjective image analysis necessitate downstream quantitative validation [29–31].

Protein and peptide arrays have enabled rapid, semi-quantitative mapping of enzyme activity and specificity in kinase and chromatin biology, accelerating motif discovery and target deconvolution [32–35]. Chemiluminescent antibody arrays extend this paradigm to cytokines and chemokines, offering quick, multiplex snapshots of pathway perturbations. However, their constrained dynamic range together with fixed panels and batch effects limit absolute quantitation and cross-study comparability, motivating orthogonal validation (e.g. ELISA, multiplex immunoassays) [29, 30]. Membrane-based cytokine arrays (e.g. Human Proteome Profiler) permit simultaneous but semi-quantitative detection across many analytes with low sample input, yet analysis often relies on subjective spot detection and heterogeneous vendor software [36, 37]. A persistent bottleneck is the scarcity of open-source, manufacturer-agnostic and automated tools for robust image processing, normalization and QC-capabilities essential for rapid, reproducible and scalable cytokine/chemokine profiling.

To address this need, we introduce InCytokine, an open-source web application for rapid relative quantification of cytokine and protein arrays. InCytokine employs image processing techniques to provide robust and reproducible data insights, shortening the cycle-time from data collection to analysis. Using InCytokine, we uncovered an intriguing expression pattern of DPP4 and ENA-78 in response to sulfatide in TREM2 variants, raising the possibility of novel interplay with TREM2 and possible implications for the role of inflammation in cardio-metabolic and Alzheimer’s diseases.

## Results

### InCytokine – a flexible framework for cytokine array analysis

A workflow describing the experimental, analytical and computational steps of InCytokine is shown in Figure 1. For illustration purposes, we used iPS-derived microglia-like (iMGL) cells treated with lipopolysaccharide (LPS) for 24 hours and cells treated with distilled water served as cognate controls. Supernatant from these treatments were incubated on a profiler array, where chemiluminescent reagents were added to generate an optical signal. A digital fluoroscopy imaging system was employed for scanning the array, resulting in the acquisition of .tiff images that were used subsequently for data processing and analysis (Figure 1A, Supplementary Figure S1-S3). Raw .tiff images were processed using the InCytokine image analysis pipeline, which operates through three automated modules. The *BlobDetector* module identifies candidate spots in array images via percentile-based contrast enhancement and multi-scale Laplacian of Gaussian filtering, followed by spatial filtering and clustering to remove duplicates and outliers. The *GridProcessor* module reconstructs the array geometry by correcting rotation, defining grid lines, and generating a reusable grid template that captures the spatial organization of all detected centroids. The *TemplateAligner* module then refines the grid-to-image correspondence through global and local optimization combined with thin-plate spline deformation, achieving sub-pixel spatial alignment. Finally, the *IntensityMeasurer* module extracts and normalizes pixel intensities within each localized spot, generating quantitative measurements for downstream analysis. The resulting log-normalized intensity tables (CSV or XLSX format) provide the basis for statistical evaluation of relative cytokine abundance and enable the identification of differentially up- or down-regulated cytokines (Figure 1B-C). InCytokine offers an intuitive user-friendly interface for scalable and reproducible image analysis of protein array scans. It features an automated, user-adjustable spot detection, template alignment, and quantitative intensity extraction, enabling reliable downstream differential abundance analysis of chemokines, cytokines and proteins from profiling assays. In addition, InCytokine backbone is modular and can be readily configured to analyze a wide variety of array-based methods.

**Figure 1.**
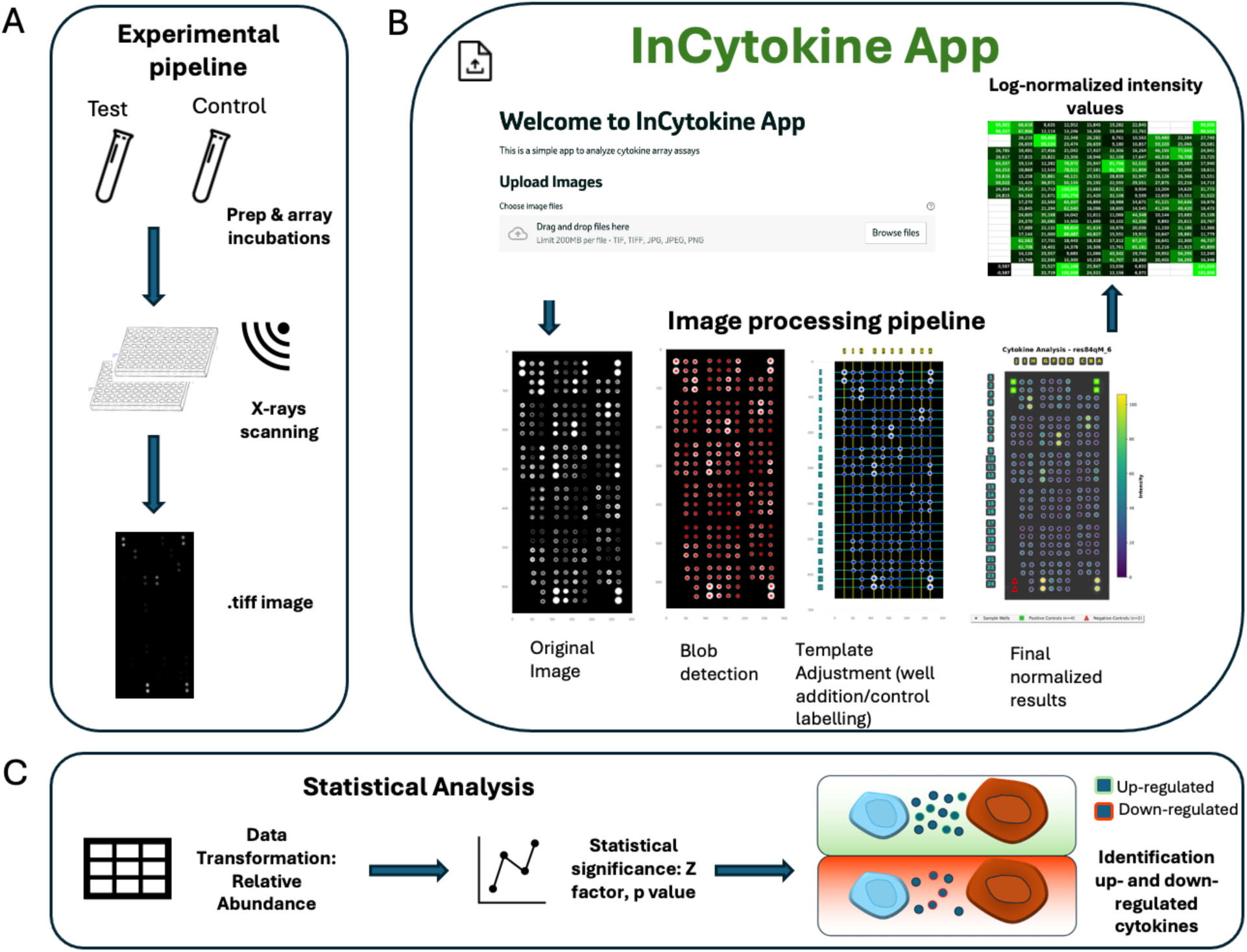
**(A)** InCytokine experimental, analytical and computational workflow (A) Treated samples (LPS/Sulfatide) and test samples (water) were prepared and incubated with the profiler array chemiluminescent reagents were added to produce optical signal. Digital fluoroscopy imaging system is used to perform X-rays and .tiff images were acquired (B) Image were processed by InCytokine pipeline through a four-stage workflow: blob detection (Laplacian-of-Gaussian with DBSCAN filtering), template generation and alignment (rotation correction, grid registration, TPS deformation), and final intensity extraction yielding log-normalized output tables. See also Supplementary Figure S1-S3 (C) InCytokine exportable files were used to compute relative cytokine abundance, and statistical tests were employed to evaluate the significance of up- and down-regulated cytokines across different conditions and genotypes.

### TREM2 mutations in iPSC-derived microglia-like cells (iMGLs) elicit differential cytokine response

TREM2 is human-genetics validated target for Alzheimer’s Disease [38–43]. Variants in TREM2 increase the risk of AD, one such being the *TREM2 R47H* variant [38–43]. We generated clonal iMGL lines with TREM2 knock-out (KO) and *TREM2 R47H* (mono-allelic clone) with wild-type (WT) iMGL lines acting as controls. These cells were treated with LPS for 24 hours and profiled them on the Proteome Profiler Human XL from R&D Systems, an ELISA-based plate capable of reporting on up to 105 cytokines, chemokines, growth factors and other soluble proteins in tissue lysates simultaneously (Figure 2A). Analyte intensity values were calculated as averages over the whole spot area and normalized with respect to the positive and negative control spots (green and red spots, respectively, Figure 2A). Fold-changes (log_2_) of the relevant protein abundances for LPS-treated analytes in *TREM2 KO*, *TREM2 R47H* point-mutation, and WT cells were computed and compared to water-treated WT cells (Figure 2B, top panels). Relative abundances were computed as the ratio between the mean abundance of the test sample and the mean abundance of the reference sample, combining four test sample values with four reference values, while only cytokines with fold-change greater than 2 in either direction (positive or negative) were considered significant (Figure 2B, top panels). The Z-factor values associated with log_2_ fold-changes were computed to determine the quality of the measurements. InCytokine provides both log_2_ fold-change and the Z-factor value for each cytokine in each treatment or genotype condition. To simplify the interpretation, both measures are used to classify the quality of the change into four distinct categories (‘ideal’, ’excellent’, ‘marginal’ and ‘inconclusive’, Figure 2B, top panels). Using a cutoff of absolute log_2_ fold-change > 1 and Z-factor > 0.5 (excellent and above), we identified IL-3 downregulation in WT cells and an upregulation of ENA-78, IL-5 and MCP-3 in *TREM2 KO* cells in response to LPS (Figure 2B, top panels). Interestingly, *TREM2 R47H* showed an increase in GROα and MCP-3 in response to LPS (Figure 2B, right-most panel & Figure 2C). The distinct set of cytokines that were altered in *TREM2 R47H* relative to *TREM2 KO* (Figure 2B, bottom panels) suggest a distinct inflammatory state for the *TREM2 R47H* cells. Interestingly, DPP4 was down-regulated in water-treated microglia cells bearing *TREM2 KO* vs *TREM2 R47H*, a role for DPP4 in TREM2 signaling or microglial biology has not been shown before.

**Figure 2.**
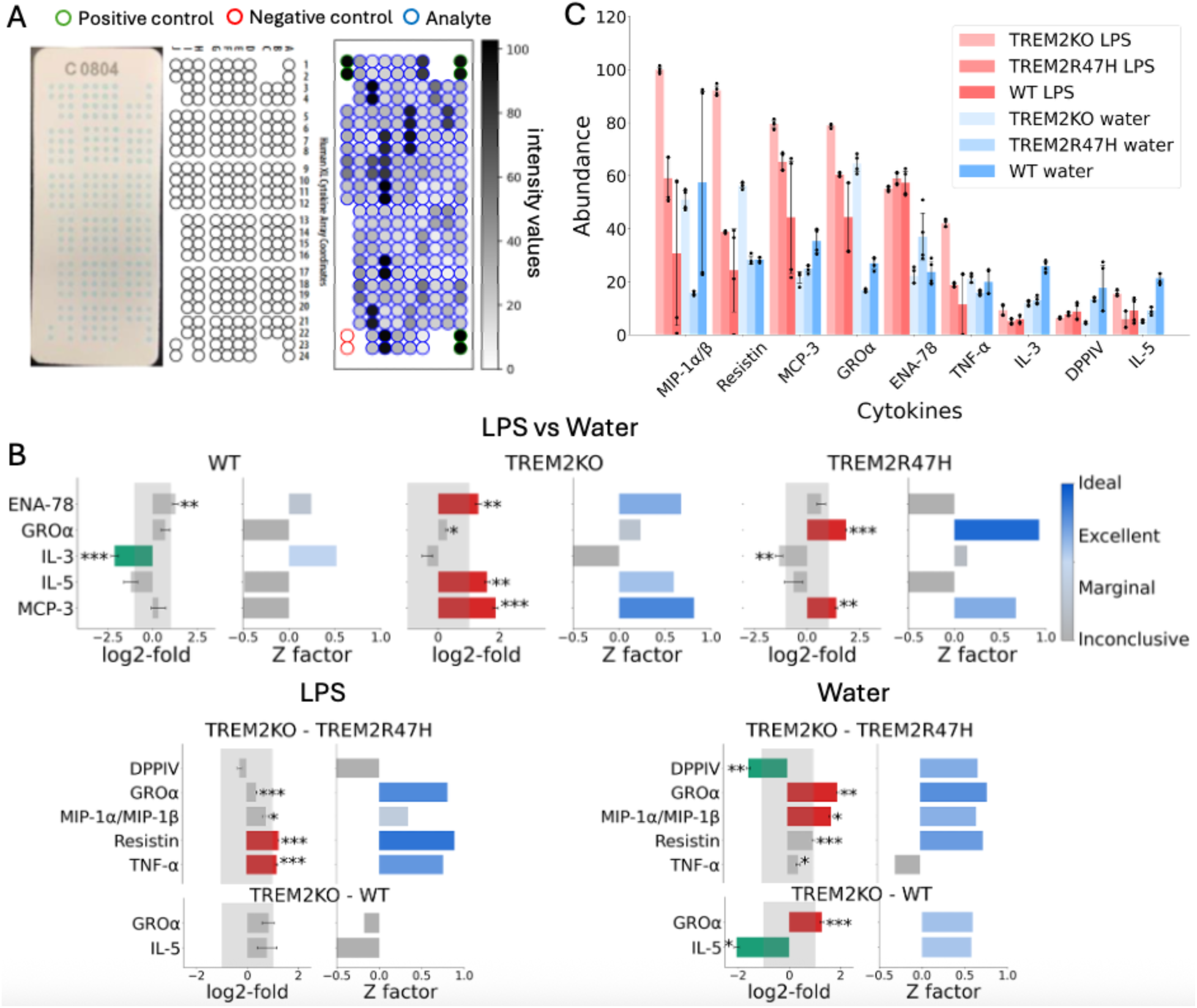
Cytokine abundance in *TREM2 KO*, *TREM2 R47H* and WT cells in response to LPS (A) Schematics of the Proteome Profiler array is an ELISA-based plate which can profile up to 105 cytokines simultaneously. On the left panel, the real array is shown; the middle panel shows the manufacturer’s schematics, while on the right panel a processed representation of the array for a single experiment (*TREM2 KO* under LPS treatment). On the corners, 3 positive and 1 negative control spots are highlighted in green and red, respectively. Analyte intensity values were calculated as average over the whole spot area, normalized with respect to the positive and negative control spots. (B) top panel: log_2_ fold-change of the relevant cytokine abundances of LPS-treated analytes versus water for *TREM2 KO*, *TREM2 R47H* allele and wild-type microglia cells. Z factor evaluates the quality of the assay measurement. Only cytokines with absolute fold-change greater than 2 with associated Z > 0.5 were considered significant. bottom panel: significant cytokines identified by comparing *TREM2 KO* and *TREM2 R47H* as well as *TREM2 KO* and WT cells under LPS (bottom left) and water (bottom right). Note that *TREM2 R47H* versus WT resulted in no significant cytokines, hence not shown C) cytokines mean intensity values of most significant cytokines (Z > 0.5, |log₂(FC)| > 1). Statistical significance was calculated using the Welch t-test p-value (only for Z > 0.5, | |log₂(FC)| > 1, *: p < 0.01, ** : p < 0.001, *** p < 0.0001).

Together, we demonstrate the utility of InCytokine as an automated tool to detect changes in the secreted proteome and uncover relevant and novel biological insights.

### Concordance between transcriptomics and cytokine abundance

Differential protein abundance can occur due to transcriptional or post-transcriptional regulation. Therefore, we next sought to reveal whether differential cytokine abundance is the direct result of transcriptional alterations. To directly test this, we performed bulk-RNA sequencing on the same cells that we used to obtain the supernatant for our cytokine experiment. Differential expression analysis of *TREM2 KO* versus *TREM2 R47H* and WT controls, and *TREM2 R47H* versus WT controls revealed a considerable number of cytokine transcripts that significantly fluctuated across treatments (LPS vs water) and/or genotype conditions (Figure 3A, p < 0.01, |log₂(FC)| > 1). Statistically significant differentially expressed transcripts that were identified across all genotype and treatment comparisons were then compared to their corresponding fold-change values as recorded on the protein arrays, showing a moderate positive correlation (Pearson’s coefficient R= 0.48, Figure 3B). Directionality remained positive even when all points with data recorded on both platforms were considered irrespective of fold-change threshold (Pearson’s coefficient R= 0.36, Figure 3B). Concordance was also apparent and significant between the sets of all genes with expression values recorded on both platforms (Fisher’s Exact test, p < 0.05, Figure 3C), as well as between the statistically significant set of transcripts detected by RNA-seq and their cytokine array counterparts (Figure 3D). A slightly better correspondence between the two technologies was found when cells were treated with sulfatide (Pearson’s coefficient R= 0.59, Supplementary Figure S4).

**Figure 3.**
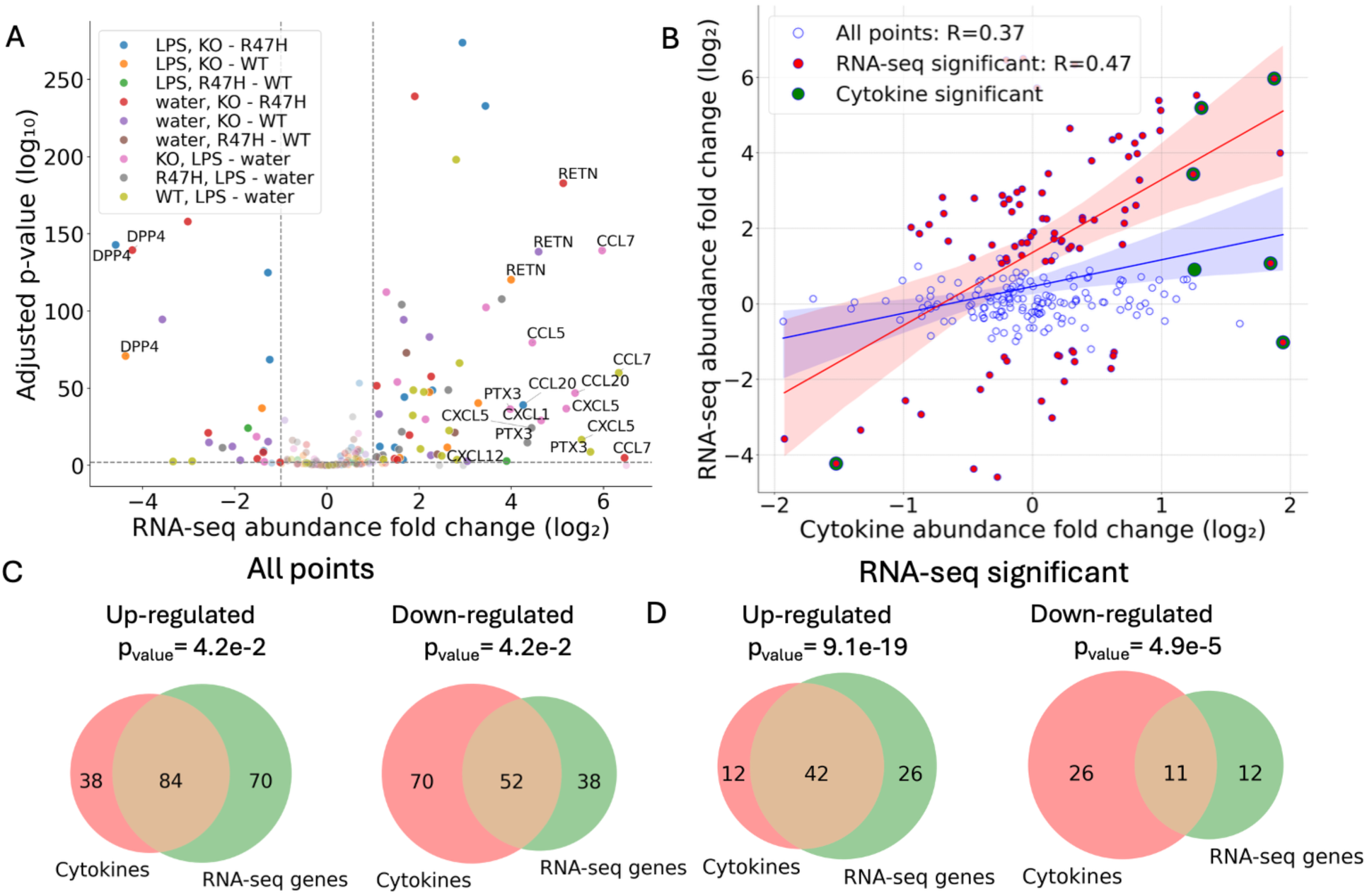
Concordance between RNA-Seq and the Proteome Profiler Human XL Cytokine Array (A) Volcano plot showing differentially expressed transcripts of all the cytokines analyzed by InCytokine; data were aggregated for all the following experiments: *TREM2 KO* vs. *TREM2 R47H* mutation (blue), *TREM2 KO* vs. WT (orange), *TREM2 R47H* mutation vs. WT (green), KO (red), *TREM2 R47H* (purple) and WT (brown) in LPS versus water. RNA-seq statistical significance was determined using adjusted p-value < 0.01 and |log₂(FC)| > 1 (B) RNA-seq correlation with cytokine relative abundance. Correlation between statistically significant transcripts (RNA-seq) and their respective cytokine abundance fold-changes (Pearson’s coefficient R= 0.48). Correlation remains positive when all points with data in both platforms were considered (Pearson’s coefficient R = 0.36). (C) The concordance of directionality of all genes and cytokines between the two platforms across all comparisons between different treatments, genotypes and controls (as shown in legend of panel A) (D) The overlap of statistically significant transcripts detected in RNA-seq with their counterparts on the cytokine array, results were aggregated for all comparisons between treatments, genotypes and controls as before. Fisher’s Exact test was used to evaluate the significance of the overlap, p-values are as indicated. See Supplementary Figure S4 for correspondence between the two platforms in sulfatide treated and untreated cells.

Despite the fundamental differences between RNA-seq and protein array-based technologies, a moderate correlation was observed between differentially expressed transcripts and their corresponding protein array values. However, there were unique examples where the abundance changes did not translate across modalities i.e. alterations in gene expression did not get reflected into cytokine-array abundance and vice versa. In such instances where differences do remain, these may represent either technical or biological effects. The latter may enhance our ability to identify transcripts for which protein expression is regulated post-transcriptionally.

### Differential response of TREM2 mutants to sulfatide

Recent studies have elucidated the complex interplay between sulfatide metabolism and TREM2-mediated microglial responses in the pathogenesis of Alzheimer’s disease and other neurodegenerative disorders. Sulfatide, sulfated galactocerebrosides highly enriched in myelin, are crucial for maintaining the structural and functional integrity of neuronal membranes and white matter. Early depletion of brain sulfatide levels is strongly associated with myelin disruptions that may amplify synaptic deficits and cognitive decline observed in AD [44, 45]. Impaired sulfatide homeostasis has been linked to enhanced neuronal vulnerability, possibly by altering lipid rafts essential for cell signaling and amyloid precursor protein processing [46]. These findings underscore the critical role of sulfatide in neuronal health and highlight the potential consequences of their dysregulation in neurodegenerative diseases.

To understand the impact of TREM2 on sulfatide metabolism, we treated WT, *TREM2 KO* or *TREM2 R47H* mutant iMGL cells with sulfatide and subjected them to cytokine array and transcriptomics profiling. In untreated cells, we noted an increase in ICAM-1 but a reduction in PDGF-AB/BB and G-CSF in *TREM2 KO* relative to WT cells (Figure 4A, top panel). However, when the same cells were treated with sulfatide, we noticed that *TREM2 KO* cells had significantly lower ICAM-1 expression along with other pro-inflammatory cytokines (Figure 4A, top panel). Lower ICAM-1 is consistent with the observed deficit in phagocytosis that is observed with *TREM2 KO* cells [47] Reduced expression of pro-inflammatory cytokines like MIP-1α/MIP-1β with a concomitant reduction in ICAM-1 mediated phagocytosis highlights how TREM2 regulates sensing and clearance of various neurotoxic agents. We further noted that DPP4, an enzyme which cleaves glucagon-like peptide-1 (GLP-1) and glucose-dependent insulinotropic polypeptide (GIP), is significantly reduced in *TREM2 KO* relative to WT in both sulfatide treated and untreated cells (Figure 4A, top panel). In contrast, *TREM2 R47H* have a distinct cytokine profile from *TREM2 KO* cells in both untreated and sulfatide-treated cells relative to WT. In untreated *TREM2 R47H* cells, cytokines and chemokines such as IL-23, MIG, MIP-3α, PDGF-AA/BB, PF4, TFF3 and TGF-α are significantly down-regulated compared to WT cells (Figure 4A, middle panel). In sulfatide treated cells, *TREM2 R47H* up-regulated ENA-78 signifying an activation of the inflammatory signaling (Figure 4A, middle panel). Finally, *TREM2 KO* cells showed higher basal levels of GROα, ICAM-1 and Resistin relative to *TREM2 R47H* in untreated cells, but substantially lower levels of these molecules in response to sulfatide (Figure 4A bottom panel). In addition, we observed lower levels of pro-inflammatory proteins such as MIP-1α/MIP-1β and ENA-78 (CXCL5) when treated with sulfatide further underscoring the inability of *TREM2 R47H* cells to respond to stimuli such as sulfatide (Figure 4A, bottom panel).

**Figure 4.**
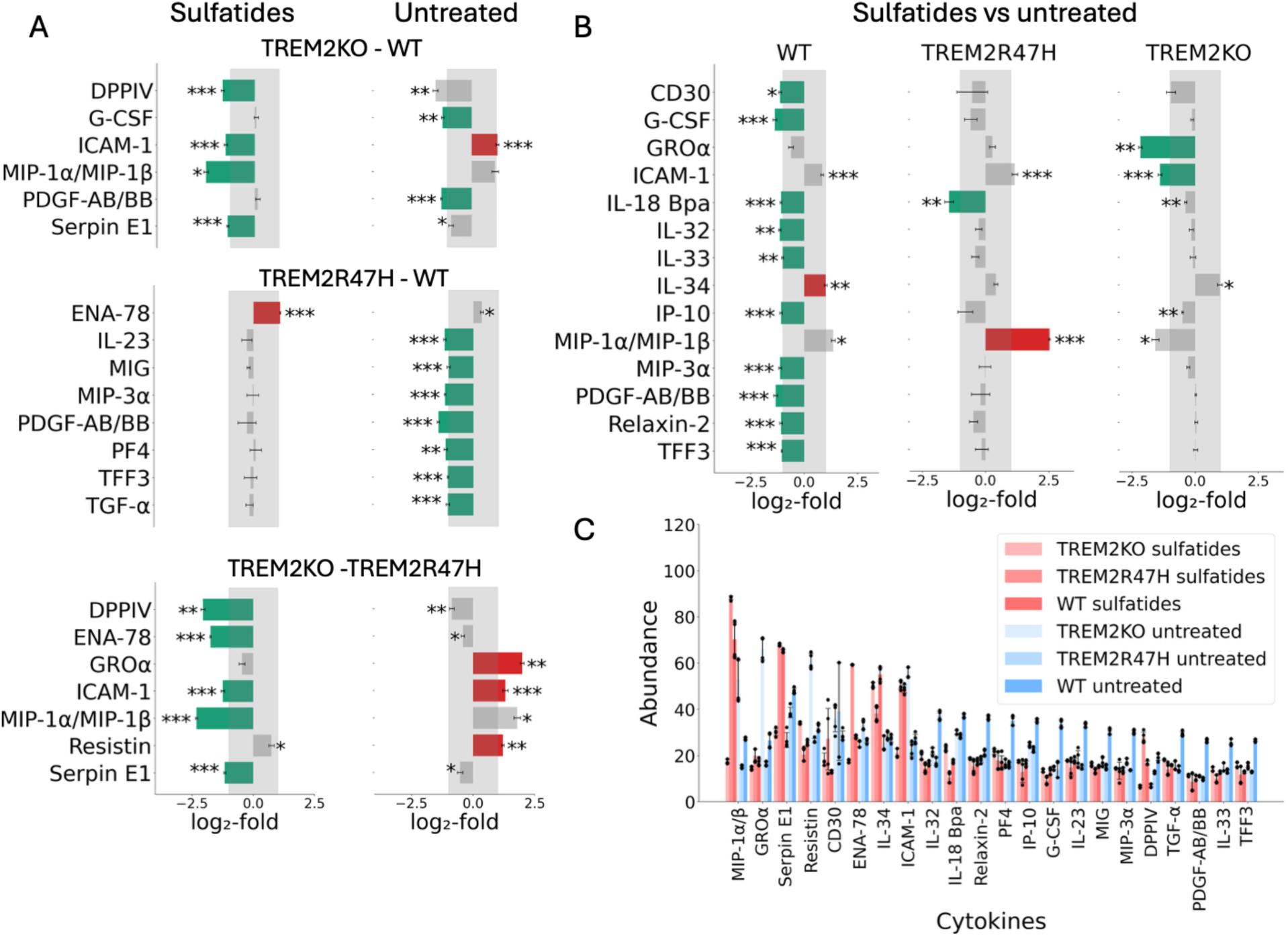
Cytokine profiling in response to sulfatide for *TREM2 KO*, *TREM2 R47H* variant and wild-type cells (A) comparison between genotypes in treated and untreated cells. Cytokine symbols are as indicated on the y-axis; log_2_ values on the x-axis; red denotes upregulation, and green down regulation; only statistically significant results are shown. Statistical significance was calculated using the Welch t-test p-value (only for Z > 0.5, |log₂(FC)| > 1,*: p < 0.01, ** : p < 0.001, *** p < 0.0001) (B) the impact of sulfatide treatment across different genotypes. Comparison between sulfatide-treated and untreated cells for *TREM2 KO*, *TREM2 R47H* and wild-type genotypes (C) Mean of the absolute intensity values of statistically significant cytokines across all samples treated and untreated.

To further parse out the sulfatide-specific impact on each genotype, we compared the response of WT, *TREM2 KO* and *TREM2 R47H* microglia to sulfatide. In addition to the genotype-specific impacts, we noted a sulfatide-specific response in WT, which was completely lacking in *TREM2 KO* and *TREM2 R47H* (Figure 4B-C). For example, WT cells down-regulated cytokines such as IL-32, IL-33, IP-10 and others, while upregulated ICAM-1 to elicit an appropriate phagocytic response and resolve inflammation (Figure 4B-C). *TREM2 KO* down-regulated ICAM-1 and *TREM2 R47H* up-regulated MIP-1α/MIP-1β in response to sulfatide (Figure 4B), underscoring inefficient phagocytosis and hyper-inflammatory response, respectively.

In summary, we uncovered a unique cytokine signature associated with *TREM2 R47H* and with *TREM2 KO* implying that the *TREM2 R47H* does not fully phenocopy TREM2 loss-of-function but provides evidence of additional functional impairments in the TREM2 pathway.

### Novel cytokines are differentially altered in response to sulfatide

Sulfatide depletions is associated with demyelination of neurons, thus amplifying synaptic deficits and resulting cognitive defects [44, 45]. Additionally, mis-regulation of sulfatide may have impacts on cell signaling pathways and amyloid precursor processing pathways [46]. One such signaling pathway which is intimately associated with Alzheimer’s disease and neuroinflammation is the TREM2 pathway. Therefore, we next sought to identify cytokines and proteins that differentially expressed in response to sulfatide stimulus and display different expression pattern in various *TREM2* genotypes.

As described in the previous section, we noticed both a sulfatide-specific and a genotype-specific response (Figure 4). We focused our attention to two analytes that showed a very distinct profile: DPP4 and ENA-78. Dipeptidyl peptidase-4 (DPP-4, CD26) is a serine exopeptidase that cleaves incretin hormones and multiple chemokines, integrating metabolic and immune signaling [43, 48]. The abundance of DPP4 across different genotypes (*TREM2 KO*, *TREM2R47H*, and WT) under various treatment conditions (sulfatide, untreated, LPS, and water) was considerably different (Table 1). While LPS treatment showed modest changes across genotype, sulfatide treatment had significantly different impacts on DPP4 levels and this was dependent on the genotype under consideration (Figure 5A). In *TREM2 KO* DPP4 (Figure 5A, blue) consistently exhibits the lowest levels across all treatments, with a log_2_ fold-change of transcripts levels ∼ -4.23 compared to *TREM2 R47H* allele in the water treatment, and ∼ -6.90 compared to *TREM2 R47H* variant in the sulfatide treatment (Table 1). DPP4 was also significantly down-regulated as compared to WT after sulfatide treatment (Table 1, log_2_ fold-change ∼ -6.05). In contrast, *TREM2 R47H* (Figure 5A, orange) showed a considerable and significant increase in DPP4 transcript level ∼2.33, particularly in response to sulfatide (Figure 5A, Table 1, Supplementary Table S1) but to a lesser extent on the protein level (log_2_ fold-change 1.13, Z-factor border significance < 0.48, Table 1), raising the possibility of an interplay between DPP4, *TREM R47H* in the context of sulfatide. Intriguingly, these patterns were captured at both the RNA and protein levels (Table 1), underscoring a unique, previously undescribed impact of TREM2 on DPP4, a key enzyme that regulates a plethora of immune-modulatory molecules.

**Figure 5.**
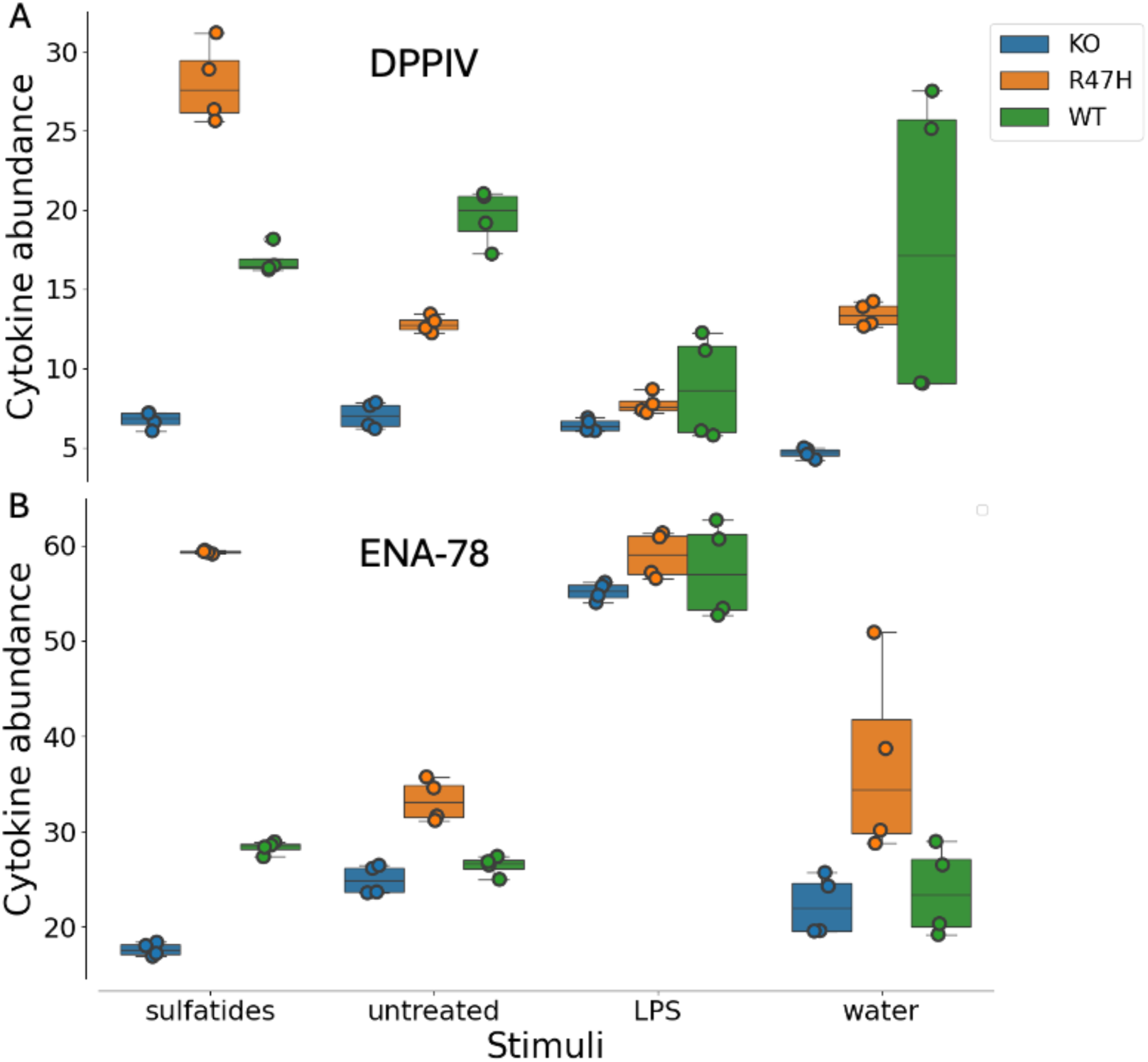
Fluctuations of DPP4 (A) and ENA-78 (B) levels in sulfatide- and LPS-treated and untreated cells for TREM2 KO (blue), *TREM2 R47H* (orange), and WT (green) genotypes. Shown on the y-axis are the mean absolute intensity values.

**Table 1.**
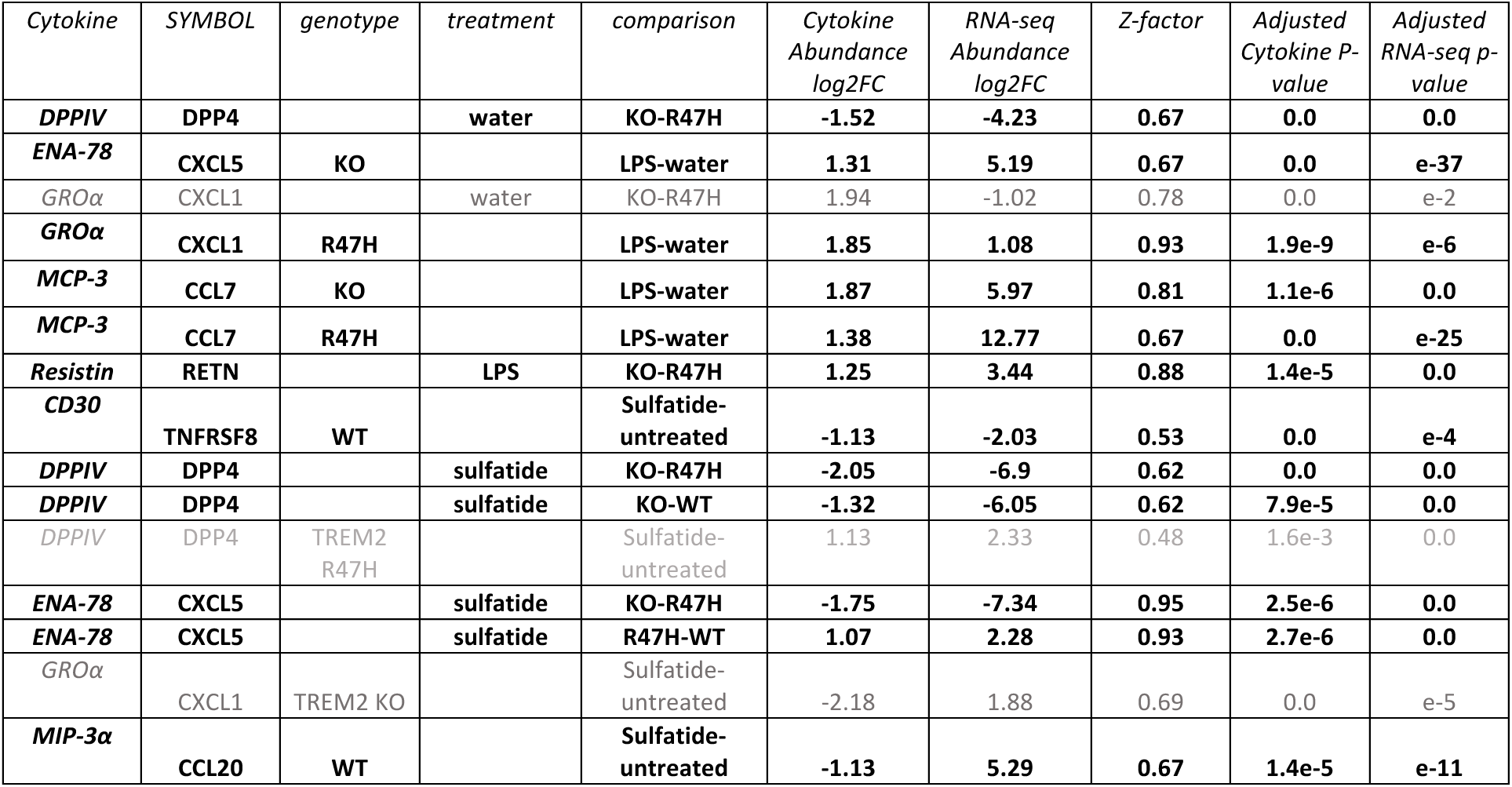
Summary table of significantly up- and down-regulated transcripts and cytokines measured by RNA-Seq and the Proteome Profiler Human XL Cytokine Array under LPS, sulfatide, and water treatments (criteria: |log₂(FC)| > 1, adjusted p-value < 0.01, Z-score > 0.5). Concordantly regulated transcript–cytokine pairs are shown in bold; discordant pairs are shaded light gray. All RNA-Seq and cytokine values across conditions and genotypes are provided in Supplementary Table S1.

| *Cytokine* | *SYMBOL* | *genotype* | *treatment* | *comparison* | *Cytokine Abundance log2FC* | *RNA-seq Abundance log2FC* | *Z-factor* | *Adjusted Cytokine P-value* | *Adjusted RNA-seq p-value* |
| --- | --- | --- | --- | --- | --- | --- | --- | --- | --- |
| **DPPIV** | **DPP4** |  | **water** | **KO-R47H** | **-1.52** | **-4.23** | **0.67** | **0.0** | **0.0** |
| **ENA-78** | **CXCL5** | **KO** |  | **LPS-water** | **1.31** | **5.19** | **0.67** | **0.0** | **e-37** |
| *GROα* | *CXCL1* |  | water | KO-R47H | 1.94 | -1.02 | 0.78 | 0.0 | e-2 |
| **GROα** | **CXCL1** | **R47H** |  | **LPS-water** | **1.85** | **1.08** | **0.93** | **1.9e-9** | **e-6** |
| **MCP-3** | **CCL7** | **KO** |  | **LPS-water** | **1.87** | **5.97** | **0.81** | **1.1e-6** | **0.0** |
| **MCP-3** | **CCL7** | **R47H** |  | **LPS-water** | **1.38** | **12.77** | **0.67** | **0.0** | **e-25** |
| **Resistin** | **RETN** |  | **LPS** | **KO-R47H** | **1.25** | **3.44** | **0.88** | **1.4e-5** | **0.0** |
| **CD30** | **TNFRSF8** | **WT** |  | **Sulfatide-untreated** | **-1.13** | **-2.03** | **0.53** | **0.0** | **e-4** |
| **DPPIV** | **DPP4** |  | **sulfatide** | **KO-R47H** | **-2.05** | **-6.9** | **0.62** | **0.0** | **0.0** |
| **DPPIV** | **DPP4** |  | **sulfatide** | **KO-WT** | **-1.32** | **-6.05** | **0.62** | **7.9e-5** | **0.0** |
| *DPPIV* | *DPP4* | *TREM2 R47H* |  | *Sulfatide-untreated* | 1.13 | 2.33 | 0.48 | 1.6e-3 | 0.0 |
| **ENA-78** | **CXCL5** |  | **sulfatide** | **KO-R47H** | **-1.75** | **-7.34** | **0.95** | **2.5e-6** | **0.0** |
| **ENA-78** | **CXCL5** |  | **sulfatide** | **R47H-WT** | **1.07** | **2.28** | **0.93** | **2.7e-6** | **0.0** |
| *GROα* | *CXCL1* | *TREM2 KO* |  | *Sulfatide-untreated* | -2.18 | 1.88 | 0.69 | 0.0 | e-5 |
| **MIP-3α** | **CCL20** | **WT** |  | **Sulfatide-untreated** | **-1.13** | **5.29** | **0.67** | **1.4e-5** | **e-11** |

A distinct expression pattern was also observed for ENA-78 chemokine, where expression level was significantly and dramatically increased in response to LPS compared to water treatment in *TREM2 KO* cells (log_2_ fold-change RNA-seq ∼ 5.19, Table 1). ENA-78 showed a significant increase in *TREM2 R47H* compared to WT (log_2_ fold-change RNA-seq ∼ 2.28, Table 1), and a dramatic decrease in *TREM2 KO* compared to *TREM2 R47H* in response to sulfatide (log_2_ fold-change RNA-seq ∼ -7.34, Figures 4B and 5B, Table 1). Epithelial neutrophil-activating peptide 78 (ENA-78/CXCL5) is an ELR+ CXC chemokine produced by epithelial, endothelial, stromal, adipose, and immune cells in response to IL-1β and TNF-α, acting primarily through CXCR2 to recruit and activate neutrophils and to influence angiogenesis and tissue remodeling [49, 50].

Together, sulfatide depletion is associated with neuronal demyelination and synaptic and cognitive deficits, and our results reveal genotype- and sulfatide-specific regulation of immune mediators. Notably a TREM2-dependent modulation of DPP4 and a divergent ENA-78 response, possibly implicating TREM2 variants in modulating inflammatory signaling relevant to Alzheimer’s disease.

## Discussion

Cytokine and chemokine profiling can shed light on the mechanisms underlying neurodegenerative diseases, and may help uncover novel therapeutic targets and, importantly, biomarkers. Here, we present InCytokine, a novel open-source tool for rapid, reproducible and scalable, semiquantitative analysis of cytokine and chemokine profiling data generated by protein array platforms. InCytokine offers an automated yet user-adjustable spot-detection and image-analysis workflow that can be readily triggered via a user-friendly interface. The output files produced by InCytokine can be used for downstream statistical analyses to identify proteins that are differentially abundant across samples and conditions.

To demonstrate the tool’s utility, we profiled TREM2 knockout, AD-associated *TREM2 R47H* variant, and wild-type cells under lipopolysaccharide and sulfatide treatments using the Proteome Profiler array. Throughout the study we applied stringent statistical thresholds to ensure biologically driven signatures over technical artifacts, accounting for measurement reliability, magnitude and the significance of the change. We observed an intriguing relationship between AD-associated risk variant in TREM2 gene and cytokine expression. These findings were also supported by transcriptomic RNA-seq data, which together suggest a potential effect of the *TREM2 R47H* variant on levels of DPP4 and other inflammatory cytokines such as ENA-78/CXCL5. We also observed good concordance between RNA-seq and Proteome Profiler despite the substantial differences in experimental and data analysis workflows.

Given differences in experimental setup and imaging equipment, imaging protocols can vary, introducing variability into the evaluation of experimental outcomes from digital array scans. InCytokine streamlines and automates the image-analysis workflow, yielding a substantial speed-up in data processing and output generation, while enforcing a standardized procedure. This enables scientists to scale up experiments and devote more time to data analysis and results interpretation. Although some manufacturers may offer software for processing digital scans, those solutions are often limited in functionality or tied to specific devices. In contrast, InCytokine is open source, allowing customization and extension while preserving the advantages of a standardized, reproducible and scalable image-processing protocol and faster experiment evaluation.

Our findings reveal that *TREM2 R47H* does not simply phenocopy TREM2 loss-of-function but rather exhibits a unique inflammatory signature, challenging the prevailing hypothesis that *TREM2 R47H* represents a simple hypomorphic allele [51, 52]. The distinct cytokine profiles observed between *TREM2 KO* and *TREM2 R47H* variants especially the differential regulation of GROα, MCP-3, and ENA-78 suggests that the *TREM2 R47H* mutation may confer both loss- and gain-of-function characteristics. This observation aligns with recent structural studies indicating that *TREM2 R47H* disrupts ligand binding while potentially altering downstream signaling in complex ways [53, 54]. The upregulation of ENA-78 (CXCL5) specifically in *TREM2 R47H* cells following sulfatide treatment represents a particularly intriguing finding. ENA-78 is a potent neutrophil chemoattractant that signals through CXCR2 and has been implicated in neuroinflammatory processes. The dramatic increase in ENA-78 in *TREM2 R47H* but not *TREM2 KO* cells in response to sulfatide suggests that this variant may promote a distinct neuroinflammatory environment that could contribute to AD pathogenesis through enhanced neutrophil recruitment and activation. This finding is consistent with recent evidence showing increased neutrophil infiltration in AD brain tissue [55, 56]. The differential sulfatide responses across TREM2 genotypes reveal critical insights into lipid sensing and myelin homeostasis. Wild-type cells appropriately upregulated ICAM-1, while suppressing inflammatory cytokines, indicative of effective phagocytic clearance with inflammation resolution. Conversely, *TREM2 KO* cells exhibited reduced ICAM-1 and inflammatory cytokine production, confirming impaired phagocytosis. Notably, *TREM2 R47H* cells showed sustained inflammation (elevated ENA-78) despite reduced phagocytic markers, uncoupling inflammation from debris clearance, a hallmark of AD pathology. Since sulfatide depletion occurs early in AD [57–59] the impaired response of TREM2 variant microglia to sulfatide-containing debris may accelerate disease progression.

Perhaps the most unexpected finding from our analysis is the identification of DPP4 as downstream of TREM2 signaling. The consistent downregulation of DPP4 in *TREM2 KO* cells contrasted with its upregulation in *TREM2 R47H* cells particularly following sulfatide treatment, suggests a previously unrecognized regulatory relationship. DPP4, also known as CD26, is a multifunctional serine protease that cleaves various chemokines and incretin hormones, thereby modulating both immune and metabolic signaling [60–62]. The differential regulation of DPP4 between TREM2 variants may have profound implications for microglial function and AD pathogenesis. DPP4 inhibition has been shown to exert neuroprotective effects in various neurodegenerative disease models [63, 64] and DPP4 inhibitors used clinically for diabetes management have been associated with reduced AD risk in epidemiological studies [65]. Our findings suggest that TREM2 status may influence the therapeutic potential of DPP4-targeted interventions, with *TREM2 R47H* carriers potentially showing different responses compared to individuals with normal TREM2 function.

## Conclusion

In conclusion, InCytokine represents a powerful tool for systematic analysis of cytokine and chemokine profiles, enabling the discovery of novel biological insights into microglial function and neuroinflammation. Our identification of distinct inflammatory signatures for TREM2 variants, the possible novel role of DPP4 in TREM2 signaling, and the differential responses to sulfatide provide important new perspectives on TREM2 biology and its role in AD pathogenesis. These findings underscore the complexity of TREM2-mediated microglial responses and highlight the need for nuanced therapeutic approaches that consider both genotype and the specific inflammatory context. Future research should focus on elucidating the precise mechanisms by which TREM2 variants influence cytokine production and exploring the therapeutic potential of modulating TREM2 signaling in AD.

## Materials and Methods

### InCytokine tool architecture

InCytokine was implemented in Python, integrating widely used open-source libraries for data science and image analysis. Numerical operations were performed with NumPy and SciPy, while image preprocessing and manipulation used Pillow (PIL) and scikit-image. Feature extraction and spatial clustering relied on scikit-learn, including KDTree-based nearest-neighbor searches and DBSCAN clustering. Tabular data handling and downstream analyses were managed with pandas, visualization with matplotlib and seaborn, and the interactive interface with Streamlit, which allows users to inspect raw images, detection overlays, intensity profiles, and processed results directly within the application. The workflow is designed to generate precise and reproducible grid-aligned spot centroids that form the spatial foundation for all subsequent intensity measurements. It operates through a structured three-stage process encompassing blob detection, grid processing, and template alignment, followed by intensity measurement and normalization.

### Image processing: grid coordinate extraction and alignment

At the first stage, the *BlobDetector* module performs a four-step detection procedure: (1) percentile-based contrast enhancement rescales image intensities (parameters: contrast_lower, contrast_upper) to improve spot visibility under varying acquisition conditions; (2) multi-scale Laplacian of Gaussian filtering identifies circular features across a configurable sigma range (min_sigma, max_sigma, num_sigma, threshold); (3) KDTree-based spatial filtering removes overlapping detections, retaining the largest centroid within a given radius (radius_filter); and (4) DBSCAN clustering eliminates isolated outliers and organizes valid centroids into a grid-like structure. This stage yields an initial geometric layout of candidate spots.

At the second stage, the *GridProcessor* module refines the geometric consistency of the detected centroids and constructs a reusable grid template. Automatic rotation correction is achieved through coordinate variance minimization, where properly aligned grids exhibit minimal Y-variance within rows and X-variance within columns. A grid search over ±30° (0.1° steps) identifies the orientation yielding the lowest within-group variance, and a rotation is applied if the confidence exceeds 0.3. Grid structure detection is then performed using independent DBSCAN clustering on X and Y coordinates to remove outliers and assign spatial indices. Linear regression through the clustered centroids defines the row and column lines, and their intersections provide precise well positions. The resulting grid structure is stored as a *GridTemplate* containing centroid coordinates, grid indices, fitted line parameters, and the median blob radius

At the third stage the *TemplateAligner* module aligns the detected centroids to the predefined grid template through a three-phase optimization strategy. Global optimization with differential evolution searches over rotation (±15°), translation (±50 px), and uniform scaling (±10%) to maximize centroid overlap without penalizing missing detections. Local refinement using the L-BFGS-B algorithm is then applied when the global solution achieves an objective score below 0.5, ensuring sub-pixel accuracy. Finally, a thin-plate spline deformation establishes point correspondences between template and detected centroids (minimum of 10 required), enabling local non-rigid warping and further improving alignment precision. The enhanced *GridTemplate* supports microplate-style well labeling (e.g., A1, B2, J23), orientation inversion, automatic remapping of control wells, and dynamic addition or removal of wells while preserving geometric consistency.

### Image processing: Intensity Measurement

Once all spots are detected and accurately aligned, the *IntensityMeasurer* module extracts and normalizes the fluorescence or colorimetric signal for each spot. Pixel intensities within each defined boundary are averaged to compute the measurement value, while control wells serve as reference points for normalization across arrays. Spurious detections are automatically excluded, and the resulting intensity matrix is exported together with visualization overlays and well labels. To maintain flexibility, each spot is represented by a two-dimensional Gaussian kernel of known diameter, and the grid is stored as a normalized coordinate template. This design allows extension to different array layouts by simply substituting the spot kernel and coordinate pattern, without altering the core detection–alignment–measurement workflow. The resulting architecture ensures that cytokine array coordinates remain consistent and reproducible across imaging conditions, providing a reliable basis for downstream quantitative analysis. These quantified values are displayed to the user through the application interface. As illustrated in Figure 1B and Supplementary Figure S1-S3, the workflow begins with the raw cytokine array image, proceeds through automated spot detection and grid alignment, and yields corrected coordinates ready for the final intensity quantification stage.

### Statistical Analysis

The log_2_ fold-change (log_2_ FC) of intensity values were obtained by comparing the base-2 logarithm of the mean absolute cytokine abundance of the 4 array analytes under either fixed treatment or genotype:

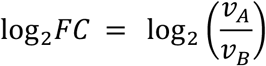

Analytes with a fold-change >=2 were considered significant (abs(log_2_*FC*) ≥ 1 for up- and down-regulated expression). The Z-factor, a statistical measure of assay quality, was used to evaluate the reliability and separation of the assay measurements:

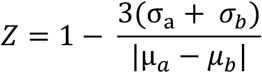

A Z-factor value greater than 0.5 was considered as a reliable measurement, while values smaller than 0.5 were considered marginal. Furthermore, a Welch t-test was conducted to evaluate the statistical significance of the observed changes, with p-values categorized as follows: p < 0.01, p < 0.001, and p < 0.0001. An RNA sequencing analysis was performed to assess concordance with cytokine array data. For RNA-seq only transcripts with adjusted P-value (p_adj_) < 0.01 were considered as statistically significant. To evaluate concordance between the two platforms (RNA-seq and protein array), results were pooled across treatments, genotypes and controls and analytes with concordant log_2_ fold-changes (i.e., directionally consistent up- or down-regulation) between RNA-seq and cytokine measurements were counted. Let N_overlap_ be the number of observations with concordant fold-change direction in both platforms (either both up or both down). Let N_cyt_ and N_RNA_ be the number of observations that are up- or down-regulated only in the cytokine dataset and in the RNA-seq dataset, respectively. We computed the following contingency table:

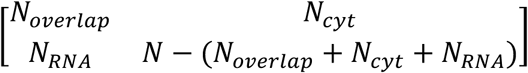

A one-sided Fisher’s exact test (alternative = “greater”) was used to assess whether the overlap count was larger than expected under the null hypothesis of independence between platforms. P-values < 0.05 were considered statistically significant.

### Sample preparation - Cytokine arrays and RNA-Seq iMGL culture

iMGLs were maintained in a complete differentiation medium prepared according to established protocols. The base medium consisted of phenol red-free DMEM/F12 (1:1), supplemented with human insulin (0.02 mg/mL), ITS-G (2% v/v), B27 (2% v/v), N2 (0.5% v/v), monothioglycerol (200 µM), GlutaMAX (1X), and NEAA (1X). This mixture was sterilized using a 0.22 µm filter. Immediately before use, the base medium was further supplemented with M-CSF (25 ng/mL) and IL-34 (100 ng/mL) to generate the complete differentiation medium. For culture, iMGLs were thawed in the base differentiation medium and centrifuged at 300 x g for 5 minutes at room temperature. After discarding the supernatant, the cells were gently resuspended in the complete differentiation medium. Cultures were incubated at 37°C in an environment containing 5% O2 and 5% CO2 for 2 to 4 days, with half of the medium replaced every 2 days.

### iMGL treatment

Six-well cell culture plates were prepared by coating each well with 1 mL of Matrigel (Corning, ref# 356231) at a concentration of 0.167 mg/mL in cold Advanced DMEM/F12 (Thermo Fisher, Cat# 12634010). The plates were incubated overnight at 4°C. The following day, iMGLs were seeded at a density of 1 × 10^6 cells per well onto Matrigel-coated plates in complete differentiation medium. Cells were cultured for 2 days at 37°C in an atmosphere of 5% O2 and 5% CO2. On the day of treatment, 0.5 mL of medium was replaced with fresh complete differentiation medium containing one of the following: water (control) or LPS (1mg/mL) or sulfatide (1mM) or left untreated. Cells were then incubated for an additional 24 hours under the same conditions (37°C, 5% O2, 5% CO2) before being collected for bulk RNA sequencing while supernatants were collected for performing cytokine arrays.

### Bulk RNA sequencing

Bulk RNA-seq was performed by Azenta Life Science. Total RNA was extracted and purified from 1,000,000 pelleted cells using TrueSeq stranded total RNA Ribo-zero kit from Illumina. For each sample, 100 ng of RNA was used to prepare the RNA library, which was then sequenced on an Illumina NovaSeq platform with a 2 × 100 bp configuration, targeting 6 Gb of data per sample. Raw FASTQ files were assessed using FastQValidate (FQV), which included adapter trimming and quality control checks. The data were then processed with Omicsoft, using the OSA Alignment algorithm and the human genome reference std b38 ensmbl r108. Aligned reads were imported into DESeq2 v1.28.1 for normalization and differential expression analysis. Genes with low abundance (total gene count < 40 across all samples) were filtered out.

### Cytokine analysis using Proteome Profiler Human XL Cytokine Array Kit

Supernatant collected from treated cells or controls were frozen at –80 °C before proceeding with cytokine analysis using the Proteome Profiler Human XL Cytokine Array Kit from the R&D Systems according to manufacturer’s instructions. Briefly, supernatant was collected after centrifuging the cells for 5 min at 1500rpm, and then 100uL of the supernatant was diluted with buffers from the Proteome profile kit followed by shaking incubation in cold room of the cytokine array overnight. Next morning, washes were performed using the wash buffers provided in the kit followed by developing the array using the near-infrared (NIR) fluorescence detection using the LI-COR Odyssey^®^ Infrared Imaging System. R&D Systems provides a modified protocol for developing these arrays using specialized IRDye^®^ 800CW Streptavidin.

#### Data availability

Bulk RNA sequencing data are available in GEO under the accession: GEO XXXXX All code for this publication along with cytokine image data are available in the following GitHub repository: https://github.com/MSDLLCpapers/InCytokine.

## Abbreviations

AD: Alzheimer’s disease
Aβ: amyloid-beta
CSV: comma-separated values
CXCL5: C-X-C motif chemokine ligand 5 (ENA-78)
DPP4: dipeptidyl peptidase-4 (CD26)
ELISA: enzyme-linked immunosorbent assay
ENA-78: epithelial neutrophil-activating peptide-78 (CXCL5)
FC: fold-change
FDR / padj: false discovery rate / adjusted p-value
GLP-1: glucagon-like peptide-1
GIP: glucose-dependent insulinotropic polypeptide
iMGL: induced pluripotent stem cell–derived microglia-like cells
JSON: JavaScript Object Notation
KDTree: k-dimensional tree (nearest-neighbor data structure)
LPS: lipopolysaccharide
NIR: near-infrared
OMIC: (used here generically) omics / high-throughput molecular profiling
PDGF: platelet-derived growth factor
QC: quality control
RNA-seq: RNA sequencing (bulk)
TREM2: triggering receptor expressed on myeloid cells 2
TIFF / .tiff: Tagged Image File Format

## Supplementary Data

Supplementary Table S1. and Figures (S1-S4) are provided.

## Acknowledgments

We thank M. Isabel Agea for assistance with backend and user interface integration, and Jaroslav Cerman for preparing the code for open-source release. We also acknowledge Dave Pace, Jyoti Shah, Antong Chen and Carol A. Rohl for supporting this work and are grateful to Michael Wurst for his invaluable manuscript review and feedback.

## Author Contributions

DB, DJ, RM and VP conceive the study. DB and DJ wrote the manuscript. AV, FO and MA also contributed to the method section of the manuscript. DJ and SF prepared the samples and performed the experiments. FO and JS developed the first version of the image analysis workflow. MV developed a prototype of the user interface. AV, IP implemented the final version of the software. AV designed and developed the automated spot detection component. MA performed all the statistical data analysis and generated the figures. All authors read, provided feedback and approved the final draft.

