## Supplementary Figure S1, Supplementary Figure S2, Supplementary Figure S3, Supplementary Figure S4 for "InCytokine, an open-source software, reveals a TREM2 variant specific cytokine signature"

### Supplementary Material

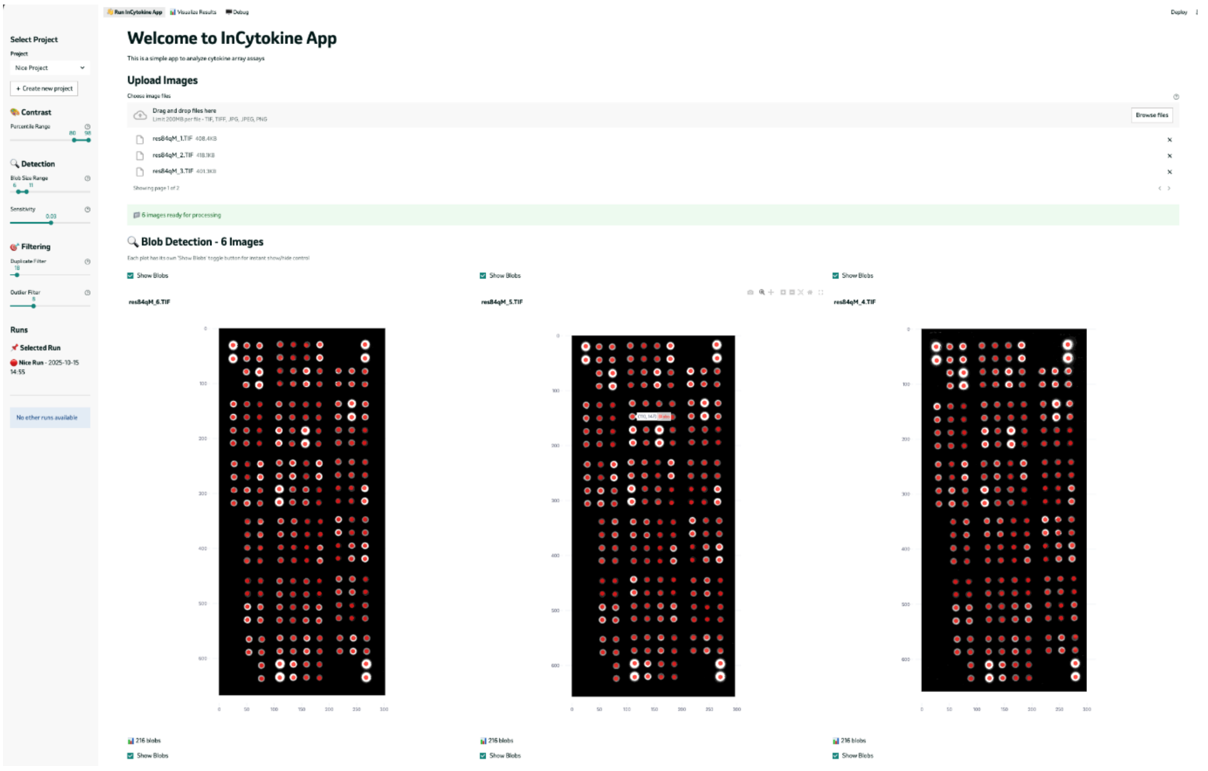


**Supplementary Figure S1.** Blob Detection module of the InCytokine App interface. The uploaded cytokine array images are processed using the multi-scale blob detection algorithm to identify well locations across all array replicates. Each detected well is outlined in red for visual validation before subsequent template alignment and intensity extraction steps.

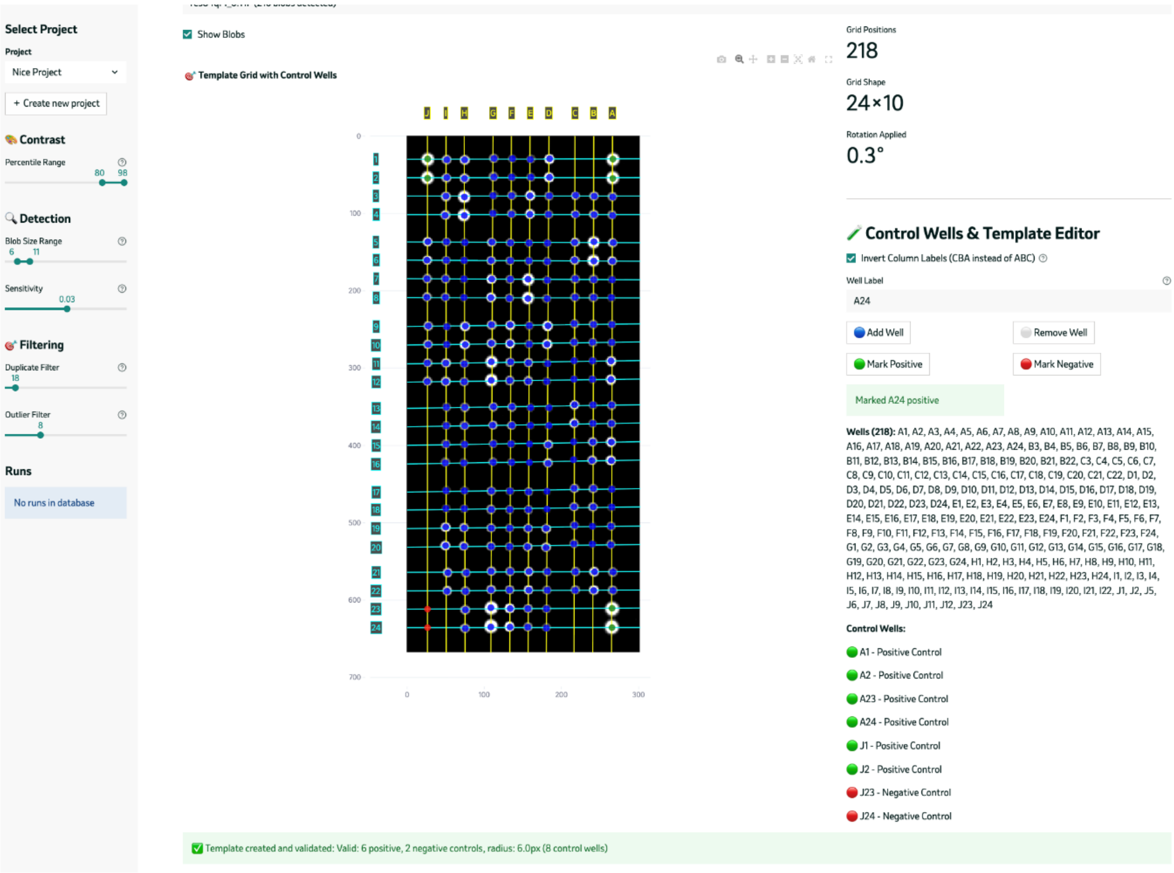


**Supplementary Figure S2.** Template Grid and Control Wells module of the InCytokine App. The detected spot array is rotated to estimate the grid lines, which enables the template adjustment (well addition/removal, control well labelling). Positive and negative control wells are interactively marked (right panel) to calibrate assay normalization and orientation. The overlay (center) displays validated grid geometry, including all detected wells (blue circles) and designated control wells (green and red markers), confirming correct template alignment before intensity quantification.

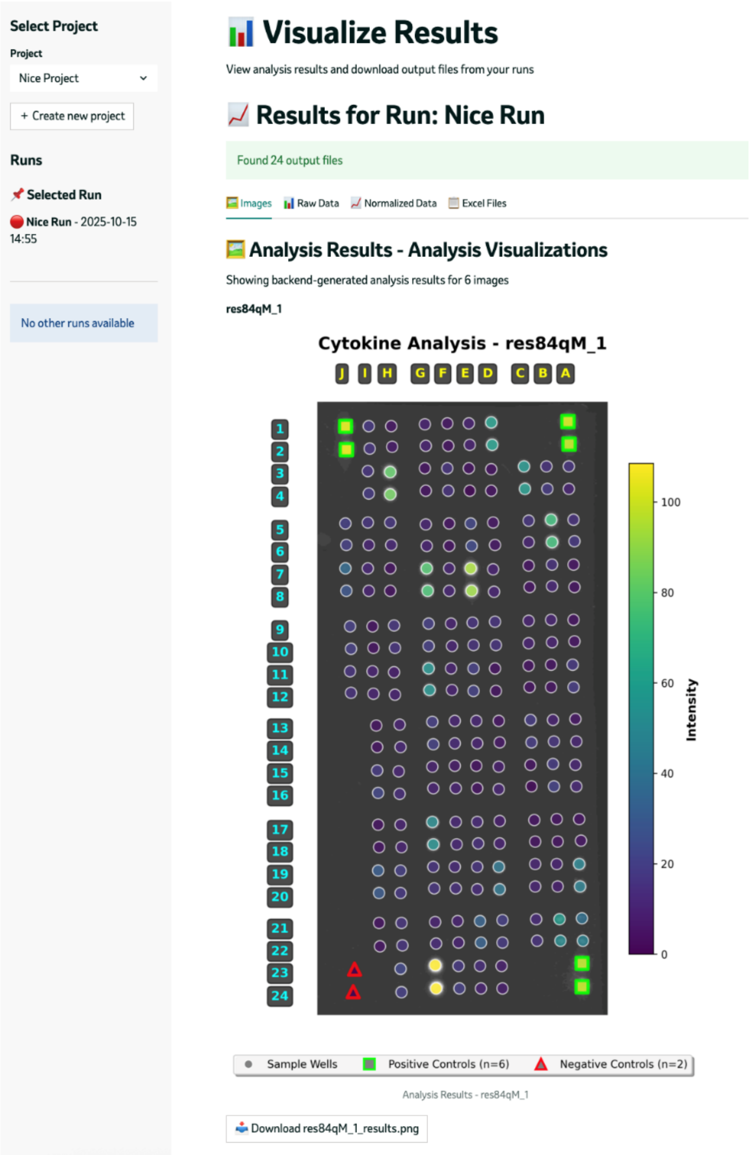


**Supplementary Figure S3.** Intensity Visualization module of the InCytokine App. Following template alignment and normalization, each cytokine's well is quantified and color-coded by measured signal intensity. Positive (green squares) and negative (red triangles) control wells define assay calibration boundaries, while sample wells (purple–cyan gradient) reflect relative cytokine abundance. The heatmap overlay enables quick visual assessment of signal distribution and assay consistency across the array.

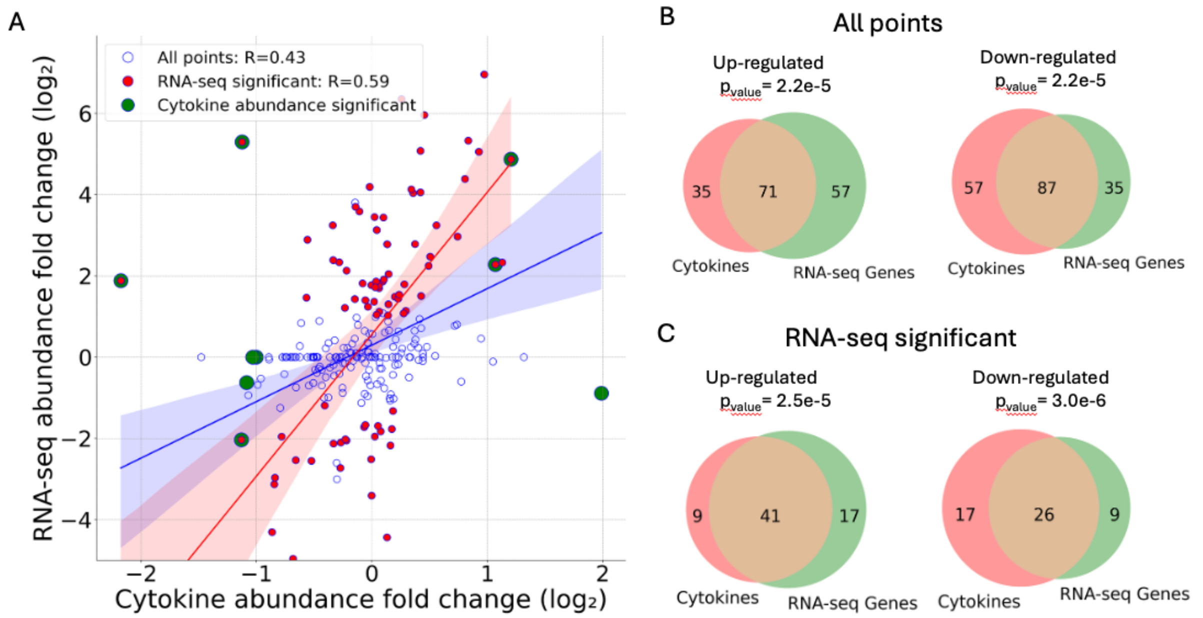


**Supplementary Figure S4.** Correspondance between RNA-Seq and ProteomeProfiler array (A) Correlation between RNA-seq and cytokine profiling. (Pearson's coefficient as indicated, R=0.49 for all points and R= 0.58 for the significant set of transcripts in RNA-seq). (B) The overlap of all transcripts with data on both platforms and (C) the overlap of statistically significant transcripts across all comparisons between treatments, genotypes and controls.
